# Phylogenetic variations in a novel family of hyperstable apple snail egg proteins: insights into structural stability and functional trends

**DOI:** 10.1101/2023.04.28.538759

**Authors:** M. Y. Pasquevich, M. S. Dreon, M. E. Diupotex-Chong, H. Heras

## Abstract

The relationship between protein stability and function evolution has not been explored in proteins from natural sources. Here, we investigate the phylogenetic differences of Perivitellin-1 (PV1) a novel family of hyperstable egg carotenoproteins crucial to the reproductive success of *Pomacea* snails, as they have evolved clade-specific protective functions.

We studied *P. patula* PV1 (PpaPV1) from Flagellata clade eggs, the most basal of *Pomacea* and compared it with PV1s orthologs from derived clades. PpaPV1 stands as the most stable, with longer unfolding half-life, resistance to detergent unfolding, and therefore higher kinetic stability than PV1s from derived clades. In fact, PpaPV1 is among the most hyperstable proteins described in nature. In addition, its spectral characteristics providing a pale egg coloration, mild lectin activity and glycan specificity are narrower than derived clades.

Our results provide evidence indicating large structural and functional changes along the evolution of the genus.

Notably, the lectin binding of PpaPV1 is less pronounced, and its glycan specificity is narrower compared to PV1s in the sister Bridgesii clade. Our findings underscore the phylogenetic disparities in terms of structural and kinetic stability, as well as defensive traits like a potent lectin activity affecting the gut morphology of potential predators within the Bridgesii clade or a conspicuous, likely warning coloration, within the Canaliculata clade.

This work provides a comprehensive comparison of the structural attributes, stability profiles, and functional roles of apple snail egg PV1s from multiple species within a phylogenetic context. Furthermore, it proposes an evolutionary hypothesis suggesting a trade-off between structural stability and the functional aspects of apple snail’s major egg defense protein.

## INTRODUCTION

Protein stability affects functional aspects and evolvability, and this has established a tight balance between increased functionality through the accumulation of mutations and, except intrinsically disordered proteins, the ability to maintain an adequate level of stability (Gershenson et al., 2014). However, there are not many examples from nature regarding these tradeoffs, as opposed to directed evolution in the laboratory (Zheng et al., 2020) or theoretical models (Agozzino & Dill, 2018; Tokuriki & Tawfik, 2009a; Zeldovich et al., 2007).

Apple snails (*Pomacea* spp.) are an emerging model for evolutionary studies due to their high diversity, ancient history, and wide geographical distribution (Hayes et al., 2009; Sun et al., 2019). *Pomacea* spp. are amphibious snails that have evolved an unusual reproductive strategy, laying eggs outside the water (Hayes et al., 2009). This transition to terrestrial egg laying went along with the acquisition of remarkable molecular and biochemical changes, particularly their reproductive egg proteins (Ip et al., 2019). Multiple lines of evidence on adaptive evolution in the egg proteins contribute to our understanding of how aquatic gastropod ancestors invaded terrestrial habitats (Ip et al., 2019).

The traditional view that proteins possess absolute functional specificity and a single, fixed structure, conflicts with their marked ability to adapt and evolve new functions and structures (Tokuriki & Tawfik, 2009b). In some animal genera, orthologous proteins play similar roles but undergo major functional adaptations. Particularly, *Pomacea* has a family of reproductive egg carotenoprotein, called Perivitellin-1 (PV1) with no sequence similarity with proteins of other organisms outside the ampullariid family, suggesting that PV1s have aroused by duplication of orphan genes. PV1s orthologous have been studied only in the most derived clades. They have, evolved different functions in different lineages while retaining other functions unchanged (Brola et al., 2020). They share several structural features: are colored, have high molecular weight oligomers, are very glycosylated, and composed of combinations of subunits with similar amino acid sequences (Brola et al. 2020). Besides, the association of PV1s with carotenoids seems to be exclusive of the *Pomacea* genus at least those from the most derived Bridgesii and Canaliculata clades (Dreon et al., 2004a; Ituarte et al., 2008; Pasquevich et al., 2014, Brola et al., 2020). Members of this novel family of invertebrate egg reproductive proteins also display high thermal and pH structural stability, as well as a kinetic stability (Pasquevich et al., 2017, Brola et al., 2020).

PV1s are massively accumulated in eggs (Giglio et al., 2016) playing a role as a storage protein and a major source of nutrients during embryo development (Heras et al., 1998). Moreover, PV1s carry and protect antioxidant carotenoids from the harsh environmental conditions of development (M. S. Dreon et al., 2004; Ituarte, Dreon, Pollero, et al., 2008; Pasquevich et al., 2014). Remarkably, they are a poor amino acids source to predators because of their low digestibility which renders them an antinutritive protein. Noteworthy, besides embryo nutrition and the antinutritive (non-digestive) role, PV1s have clade-related functions according to their phylogenetic position: those PV1s from the Canaliculata clade, PcOvo, and PmPV1, provide a bright reddish coloration possibly a warning signal, an ecological function that would go along with the presence of a toxic perivitellin, PV2 only found in the Canaliculata clade (Giglio et al., 2020; Heras et al., 2008). On the contrary, the members of the Bridgesii clade lay more pale eggs of pinkish color (presumably non-warning signal) and have PV1s like PsSC from *P. scalaris* (Ituarte et al., 2008, 2010, 2012) and PdPV1 from *P. diffusa* (Brola et al., 2020) that possess a strong lectin activity, *i. e.* capacity to recognize and bind glycans, and adversely affect gut morphophysiology of predators, a role absent in PV1s from the Canaliculata clade. These different functions among orthologous PV1s proteins (Brola et al. 2020) go along with their remarkably high stability and provide a unique and unexplored model to understand the evolution of hyperstable proteins, an aspect poorly studied experimentally in biological models.

We began by studying the structure, stability, and functional features of PV1s from a species of Flagellata the most basal clade of the genus, which allowed a phylogenetical analysis of the clade-related structural and functional trends of these fascinating hyperstable proteins. We found that variations in this novel family of reproductive proteins accompanied the diversification of the genus and may have facilitated some apple snails to become notorious invasive pests.

## MATERIAL AND METHODS

### Sample collection and PV1s purification

*Pomacea patula* eggs were collected in the Catemaco Lake, Veracruz, Mexico, and kept in the laboratory at -70 □C until processed. The perivitelline fluid (PVF) was obtained as previously described (Pasquevich et al., 2014). In short, eggs were homogenized on ice in Tris/HCl 20 mM buffer 1:3 w/v and sequentially centrifuged at 4 □C for 20 min. at 10.000 xg and 50 min at 100.000 xg. The obtained supernatant contains the soluble egg fraction.

To compare and trace the evolution of PV1 carotenoproteins from *Pomacea*, PV1 from *P. patula catemaquensis* (hereafter PpaPV1) was purified from egg clutches as previously described for others apple snail eggs carotenoproteins (see below). Total protein was quantified following the method described by Bradford using bovine serum albumin (BSA, Sigma cat. 7906) as standard. Purity was checked by polyacrylamide gel electrophoresis (PAGE). Another *Pomacea* spp. carotenoproteins (*i. e.* PcOvo, PmPV1, and PsSC), used in some assays, were purified as previously described (Dreon et al., 2004b, Ituarte et al., 2008, Pasquevich et al., 2014).

### Anti-PpaPV1 rabbit serum preparation

Antibodies directed against purified PpaPV1 were prepared in rabbits. Animals were given a first subcutaneous injection of 120 μg of PpaPv1 emulsified in Freund’s complete adjuvant (Sigma Chemicals, St. Louis, MO, USA). A booster injection with about 60 μg antigen mixed with Freund’s in-complete adjuvant was administered after 2 and 4 weeks. Two weeks later, the rabbits were bled through cardiac puncture. The collected blood was allowed to clot overnight (4°C) and after centrifugation the serum obtained was stored at –70°C, and used in the Western Blot technique. The specificity of the rabbit antiserum against PpaPV1 was verified by immunoblotting PpaPV1 with a non-immunized rabbit serum.

### Structure

#### Oligomer and subunits electrophoretic behavior

Gel electrophoresis was used to characterize and compare the oligomer and subunits of PpaPV1 with other *Pomacea* PV1s.

Native PAGE with Laemmli buffer (pH 8.8) without SDS was performed in 4–20% gradient polyacrylamide gels in a miniVE Electrophoresis System (GE Healthcare, Life Science). High molecular weight standards (Pharmacia) were run in the same gels. Subunits were separated by SDS-PAGE in 4–20% gradient polyacrylamide gels containing 0.1% SDS; samples were denatured at 95 °C, with dithiothreitol and β-mercaptoethanol treatment (Laemmli, 1970). Low molecular weight standards (Pharmacia) were used, and gels were stained with Coomassie Brilliant Blue R-250 (Sigma Chemicals). In both gels, PcOvo PmPV1 and PsSC were run for comparison of PV1s from other clades.

#### Immunoblotting

Cross-reactivity of PpaPV1 with PV1s from other clades was assayed with anti-PpaPV1, anti-PsSC and anti-PcOvo sera. PV1s (7 μg) and molecular weight marker (Dual Color, Bio-Rad) were transferred from SDS-PAGE 16% gels onto nitrocellulose membranes (Amersham) in a Mini Transblot Cell (Bio RadLaboratories, Inc.), using 25 mM Tris–HCl, 192 mM glycine, 20% (v/v) methanol, pH 8.3 buffer. After blocking for 2 h at room temperature with 3% (w/v) nonfat dry milk in PBS-Tween, the membranes were incubated overnight at 4 °C with polyclonal antibodies against PcOvo (Dreon et al., 2003) in 1:10.000 dilution, polyclonal antibodies against PsSC (Ituarte et al., 2008) in a 1:12.000 dilution, and polyclonal antibodies against PpaPv1 (1:1000) in 3% (w/v) nonfat dry milk in PBS-Tween. Specific antigens were detected after incubating 2 h at room temperature goat anti-rabbit IgG horseradish peroxidase conjugate (BioRad Laboratories, Inc.). Immunoreactivity was visualized by electro-chemiluminescence in a Chemi-Doc MP Imaging System (Bio Rad).

#### Size Exclusion Chromatography

Size exclusion chromatography (SEC) allows us to estimate the molecular weight of proteins comparing the chromatographic retention times with those of standard proteins of precise weight (Barth & Boyes, 1990). The molecular weight of PV1s by SEC was analyzed with an SEC-Superdex 200 10/300 GL column (Amersham) in an isocratic size exclusion HPLC (1260 Infinity, Agilent technologies) with UV detection in isocratic mode. The Mobil phase contained 137 mM. NaCl, 2.7 mM KCl, 2.7 mM KCl mM, 10 mM, Na_2_HPO_4_, 1.8 mM KH_2_PO_4_ (PBS). The flow rate was 0.5 mL/min and the detector was set at 280 nm. The elution volume (Ve) of PV1s and 500 μL of standard proteins in PBS (5 mg/mL Thyroglobulin (MW 669000), 2.8 mg/mL Ferritin (MW 440000), 4 mg/mL Aldolase (MW 159000) and Ovoalbumin (MW 45000)) were determined by measuring the volume of the eluent from point of injection to the center of the elution peak. Blue Dextran 2000, 1 mg/mL was used to calculate the column void volume (V_0_). The calibration curve was performed by fitting a curve in a plot of K_av_ = (V_e_-V_0_)/(V_c_-V_0_), where V_c_ is the geometric volume of the column (24 mL), versus the log molecular weight of each standard. A standard curve was fitted to a line and PV1s MW calculate extrapolating from the standard curve.

#### Dynamic Light Scattering

The dynamic light scattering (DLS) of a nanoparticle sample in solution revealed the particle size distribution in real-time (Falke & Betzel, 2019). DLS is particularly sensitive to large aggregates, common in some PV1s (Ituarte et al., 2008, Brola et al., 2020). Thus, DLS analysis was performed in line right after SEC-eluted PV1s. PV1s sizes were monitored in PBS at 25°C, using a Malvern Zetasizer nano-zs instrument. The scatter light signals were collected at a 173-degree scattering angle (backscatter) and three measurements of an automatic number of runs each were conducted per sample. Protein parameters were analyzed with the Zetasizer Software v 7.13. Data used for size measurement meets quality criteria. Intensity size distributions were used to size calculation. Volume size distribution was used to check the main peak contribution to the analysis.

#### N-terminal sequence

Subunits of purified PpaPV1 separated by electrophoresis were transferred to PVDF membranes and sequenced by Edman degradation at the Laboratorio Nacional de Investigación y Servicios en Péptidos y Proteínas (LANAIS-PRO, Universidad de Buenos Aires—CONICET). The system used was an Applied Biosystems 477a Protein/Peptide Sequencer interfaced with an HPLC 120 for one-line phenylthiohydantoin amino acid analysis. N-terminal sequences were compared with other *Pomacea* sequences using the multiple sequence alignment program CLUSTAL 2.1 (Larkin et al., 2007).

#### Spectrophotometric analysis

Absorption spectra of egg carotenoproteins are valuable taxonomic characters in *Pomacea* spp. (Pasquevich & Heras, 2020). PV1s absorb light in the visible region of the spectrum because of the presence of carotenoid pigments (Heras et al., 2007). Absorption spectra of PVF and purified carotenoproteins were recorded between 350 nm to 650 nm in an Agilent 8453 UV/Vis diode array spectrophotometer (Agilent Technologies, Waldbronn, Germany).

## STRUCTURAL AND KINETIC STABILITY

### Effect of pH and temperature on structural stability

To study the effect of pH on PpaPV1 structural stability, the protein was incubated overnight in different buffers ranging from pH 2.0 to 12.0 following a previously used method (Pasquevich et al., 2017). Samples were analysed by absorbance and fluorescence spectroscopy. Absorbance spectra were recorded between 300–650 nm in an Agilent 8453 UV/Vis diode array spectrophotometer (Agilent Technologies, Waldbronn, Germany) taking advantage of the fact that PV1s absorb in the visible range allowing to follow the protein-carotenoid interaction by its spectrum in this range. Fluorescence emission was recorded as described in the *Chemical denaturation* section (see below). Two independent samples were measured, and the corresponding buffer blank was subtracted. The effect of temperature on PpaPV1 at pH=7.4 was also measured by absorption and fluorescence spectroscopy in the range of 25–85 °C. The effect of extreme thermal conditions was analyzed by boiling PpaPV1 for 50 min evaluating the oligomer integrity using native (non-denaturing) gel electrophoresis, as previously done (Pasquevich et al. 2017).

### Chemical denaturation

The intrinsic fluorescence emission of PpaPV1 and PsSC tryptophans was used to follow the PV1s denaturation induced by guanidine hydrochloride (GdnHCl) (Sigma). Chemical denaturation was performed by incubating overnight 50 μg/mL of PV1s in the presence of increasing concentrations (0–6.5 M) of GdnHCl buffered with 0.1 M phosphate buffer at pH 7.4 at 8°C.

Protein intrinsic fluorescence spectra were recorded on a Fluorolog 3 Spectrofluorometer coupled with a Lauda Alpha RA 8 thermostatic bath. Fluorescence spectra were recorded in emission scanning mode at 25 °C. Tryptophan emission was excited at 295 nm (6 nm slit) and recorded between 310 and 450 nm (3 nm slit). The corresponding buffer blank was subtracted. Two independent samples were measured. Spectra were characterized by their center of mass (CM) and the populations associated with the unfolded fraction (ƒu) were calculated from the CM as for PmPV1 in Pasquevich *et al*. (2017). The equilibrium reached in each GdnHCl concentration allowed the calculation of an equilibrium constant K=ƒu/(1-ƒu) and Gibb’s free energy for the unfolding reaction in terms of this mole fractions (ΔG^0^= −RT lnK) were calculated. The dependence of ΔG^0^ on GdnHCl concentration can be approximated by the linear equation ΔG^0^ =ΔG^0^H_2_O – *m*[GdnHCl], where the free energy of unfolding in the absence of denaturant (ΔG^0^H_2_O) represents the conformational stability of the protein. The GdnHCl concentration in which half of the protein is unfolded (C_m_) was estimated as a function of denaturant concentration from the linear extrapolation method.

### Resistance to sodium dodecyl sulfate

Resistance to sodium dodecyl sulfate (SDS)-induced denaturation serves to identify proteins whose native conformations are kinetically trapped in a specific conformation because of an unusually high-unfolding barrier that results in very slow unfolding rates. The resistance to SDS was assayed following the Manning and Colon procedure previously used with other *Pomacea* spp. carotenoproteins (Pasquevich et al 2017, Brola et al. 2020). Briefly, PcOvo, PmPV1, PsSC, and PpaPV1 were incubated in Laemmli sample buffer (pH=6.8) containing 1% SDS and either boiled for 10□min or unheated before its analysis by 4–20% SDS-PAGE. The gels were then stained with Coomassie blue.

### Unfolding Kinetics induced by GdnHCl

Unfolding of proteins in increased concentrations of GdnHCl allows us to calculate the rate of unfolding (half-life) in the absence of denaturant under native conditions (Manning & Colón, 2004a). A fluorolog-3 (Horiba□Jobin Yvon) fluorometer was used to measure the PpaPV1, PsSC, and PmPV1 kinetics of unfolding. Protein solutions in 100 mM phosphate buffer, pH 7.4 (PB) were treated with GdnHCl solution in the same buffer. For the fluorescence kinetic experiment, increasing concentrations of GdnHCl in PB in a 10 mm pathlength cuvette were incubated. PV1s (final concentration of 50 µg/mL) were mixed by pipetting up and down with a 1 mL pipette. Data collection was started after the chamber was closed. The shutters open automatically. Time zero was manually determined as 10 sec after the protein was added to the denaturant. The excitation/emission was 299/360 nm. The relaxation time was fit to an exponential equation. The unfolding constants were obtained for each GdnHCl concentration. The rate constants as a function of GdnHCl were extrapolated to native conditions to obtain an estimate of the rate constant (*k*) in the absence of a denaturant. Half-life was calculated as ln 2/*k*.

### Resistance to Proteolysis: proteinase K assay

Protein structural rigidity makes proteins resistant to proteolysis. PV1s rigidity was assayed by Proteinase K treatment, performed following Kim *et al*. (2004) using the concentrations modified by Frassa *et al*. (2010) and previously performed to PmPV1 (Pasquevich et al., 2017). PpaPV1 (1 mg/mL) was incubated with proteinase K (1, 10, and 100 μg/mL) in 50 mM Tris/HCl buffer (pH 8.0) containing 10 mM CaCl2 at 37 °C for 30 min. Digestion was ended by boiling samples in SDS sample buffer, and products were analysed by SDS-PAGE as above.

### Functions of PpaPV1 Hemagglutinating activity

PV1s of the Bridgesii clade has a strong lectin activity and ability to agglutinate rabbit erythrocytes (Ituarte et al., 2012, Brola 2020), but PV1s of canaliculata clade lack this capacity (Pasquevich et al., 2017). We tested this capacity in *P. patula* PpaPV1 using the same methodology. In short, PpaPV1 hemagglutinating activity was tested by hemagglutination of red blood cells (RBC) from rabbits obtained in facilities of the University of La Plata. Erythrocytes were prepared as stated in Ituarte et al. (2012) with minimal modifications. Two-fold serial dilutions of PpaPV1 in PBS (25 µl) were incubated with an equal volume of 2% (v/v) erythrocytes in PBS in U-shaped microtiter plates (Greiner Bio-One) at 37°C for 2 h. The initial PVF protein concentration was 6.7 mg/mL and the PpaPV1 concentration was 1.3 mg/mL. Results are expressed as the lowest protein concentration showing visible hemagglutinating activity by the naked eye.

### Specificity for glycan-binding

Glycan binding specificity of PpaPV1 was determined at the Core H of the Consortium for Functional Glycomics (http://www.functionalglycomics.org, Emory University, Atlanta, GA, USA). To detect the primary binding of PpaPV1 to glycans, the protein was fluorescently labeled using the Alexa Fluor 488 Protein Labeling kit (Invitrogen, Life Technologies-Molecular Probes) according to the manufacturer’s instructions. Protein concentration and the degree of labeling were determined spectrophotometrically. Fluorescently labeled PpaPV1 was assayed on a glycan array that comprised 585 glycan targets (version 5.4). The highest and lowest points from a set of six replicates were removed and the four remaining values were averaged. PpaPV1 glycan microarray data were compared with PsSC (Ituarte et al., 2018) and PdPV1 (Brola et al., 2020) microarrays using the Glycan Array Dashboard (glycotoolkit.com/GLAD/) (Mehta & Cummings, 2019).

### *In vitro* intestinal digestion and high proteolysis assays

The simulated gastroduodenal digestion of PpaPV1 was analyzed by sequentially incubating the protein for 2 h with pepsin (gastric) and 2 h with trypsin (intestinal) at 37 °C, using the method described by Moreno *et al*. (2005), with some modifications as described in Pasquevich et al. 2017. Briefly, PpaPV1 in double-distilled water was dissolved in simulated gastric fluid (SGF) (0.15 M NaCl, pH 2.5) to a final concentration of 0.5 μg/μL. Digestion commenced by adding porcine pepsin (Sigma, cat. P6887) at an enzyme: substrate ratio of 1:20 (w/w). Gastric digestion was conducted at 37 °C with shaking for 120 min. Aliquots of 5 μg protein were taken at 0, 60, and 120 min for SDS-PAGE. The reaction was stopped by increasing the pH with 150 mM Tris/Cl buffer pH 8.5. Samples were immediately boiled for 5 min in SDS electrophoresis buffer with β-mercaptoethanol (4%) and analysed as described above.

For *in vitro* duodenal digestion, the gastric digest was used as starting material. The digest was adjusted to 8.5 and sodium taurocholate (Sigma) was added. The simulated duodenal digestion was conducted at 37 °C with shaking using bovine pancreas trypsin (Sigma cat. T9935) at an enzyme: substrate ratio of 1:2.8 (w/w). Aliquots were taken at 0, 60, and 120 min for SDS-PAGE analysis. BSA was used as positive (with enzyme) and negative (without enzyme) control in both gastric and duodenal digestion.

## RESULTS

### STRUCTURAL FEATURES

SEC analysis indicate that native PpaPV1 is 265.9 kDa which is rather similar to that of all other PV1s (**Table 1**). DLS analysis showed a single main peak of the distribution of intensity and volume in the 12.2-14.0 diameter size range for PV1s from the 3 clades (**Fig. 1A, Fig S1**). PV1s from basal clades also showed a minor aggregation peak (**Fig. S1B**).

**Figure 1.**
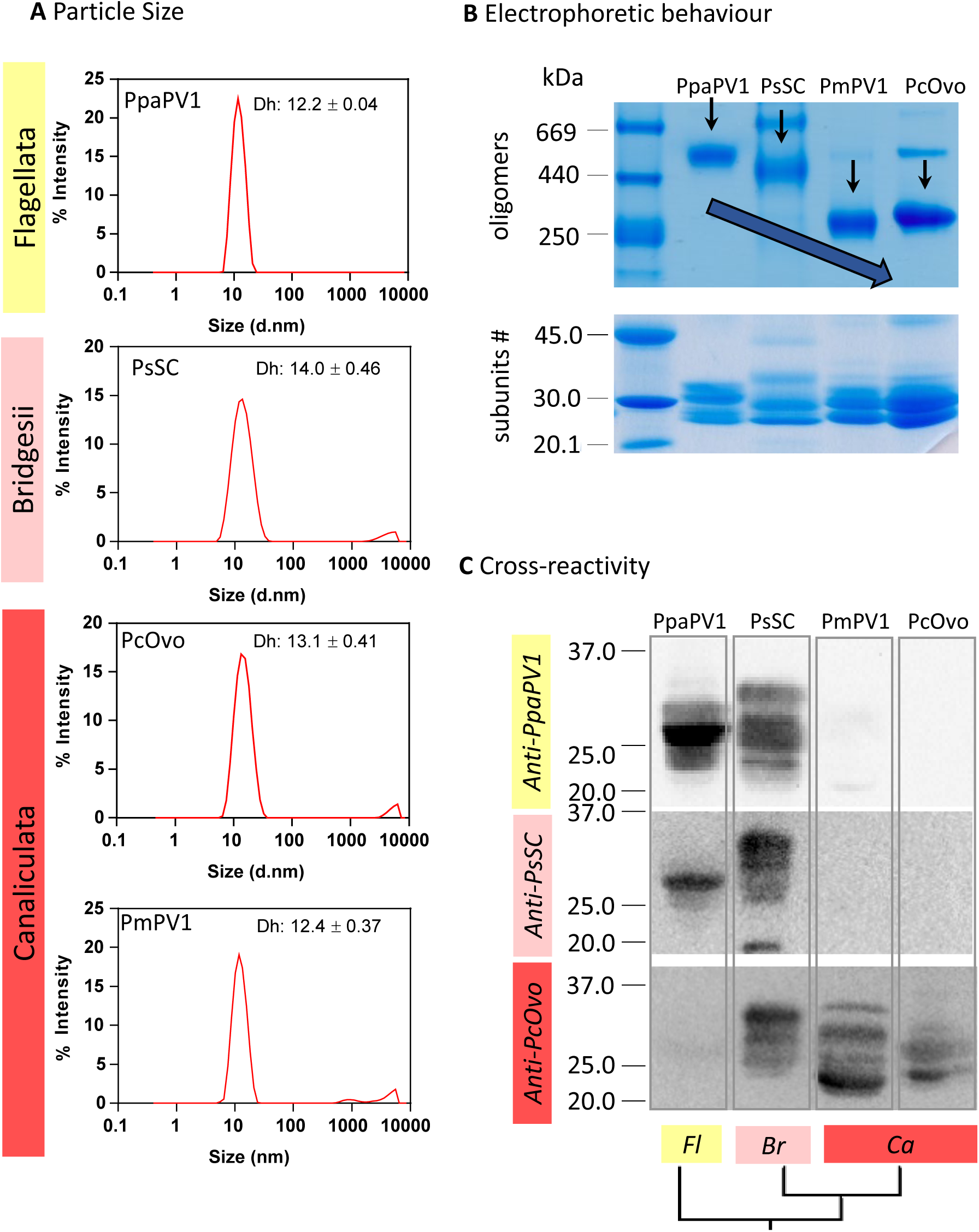
Apple snail egg PV1s have similar size and number of subunits, but different electrophoretic migration, charge surface, and immune cross-reactivity in a clade-related fashion. A. Particle size analysis as determined by DLS indicates that all PV1s have similar size and MW. B. Oligomers of PV1s in a native PAGGE (top panel). Arrows indicate the relative mobility of PV1s. The blue arrow highlights the increase in mobility along clades. Subunits of PV1s in an SDS-PAGE (lower panel). C. Western blot analysis using anti-PpaPV1, anti-PsSC, and anti-PcOvo sera. Line 1, molecular weight marker. Line 2, PpaPV1. Line 3, PsSC. Line 4, PmPV1. Line 5, PcOvo. Fl: Flagellata; Br: Bridgesii; Ca: Canaliculata. PV1 was purified from: PpaPV1 from P. patula; PsSC from P. scalaris; PmPV1 from P. maculata; PcOvo from P. canaliculata.

**Table 1.** Size and molecular weight estimation of PV1s.

|  | MW (kDa) | Diameter Size (nm) |
| --- | --- | --- |
| PpaPV1 | 267.5 | 12.2 |
| PsSC | 270.3 | 13.0 |
| PcOvo | 269.8 | 13.4 |
| PmPV1 | 265.6 | 12.4 |

Electrophoretic analysis revealed that under native conditions the mobility of PpaPV1 differs from other PV1s of the derived clades: PsSC from the Bridgesii clade, PcOvo and PmPV1 from the Canaliculata clade (**Fig. 1B**). Under denaturing conditions (SDS-PAGE) PpaPV1 showed several subunits between 25 and 35 KDa, the same molecular weight range as reported for the other PV1 of the genus (**Fig. 1B**).

Both anti-serum against PpaPV1 (Flagellata clade) and PsSC (Bridgesii clade) cross-reacts with PsSC (Bridgesii clade) and PpaPV1 (Flagellata) respectively, but not with PmPV1 and PcOvo (Canaliculata clade) **(Fig. 1C)** while PcOvo anti-serum recognized Canaliculata and Bridgesii clades PV1s (*i.e.* PcOvo, PmPV1, and PsSC) as previously described (Dreon et al., 2003, Pasquevich et al., 2017, Brola et al., 2020) but not subunits of PpaPV1 (**Fig. 1C**).

PpaPV1 was separated into 6 subunits numbered according to their electrophoretic mobility. The seven N terminal sequences obtained were grouped into two nearly identical sequences (**Fig. S2-A**). Sequence similarity analysis with PcOvo, PmPV1, and PsSC. PpaPV1_3a revealed 70.6% similarity with all three PV1s (**Fig. S2-B**), while PpaPV1_6 presents 70.6% similarity with PsSC (Bridgesii clade) and 55.6% with PmPV1 and PcOvo (Canaliculata clade) (**Fig. S2-C**). PpaPV1_3b Nt region has no sequence similarity with any reported *Pomacea* spp. perivitellin.

*P. patula* PpaPV1 and PVF absorb in a wide range of the visible spectra (350-650 nm) (**Fig. 2**) with a maximum at 380 nm an absorption maximum shared with Bridgesii clade (*P. scalaris*) but that differs from Canaliculata clade PV1s. Besides, the overall absorption intensity between 450-600 nm increased according to the phylogenetic position from Flagellata towards Canaliculata clades (**Fig. 2**).

**Figure 2.**
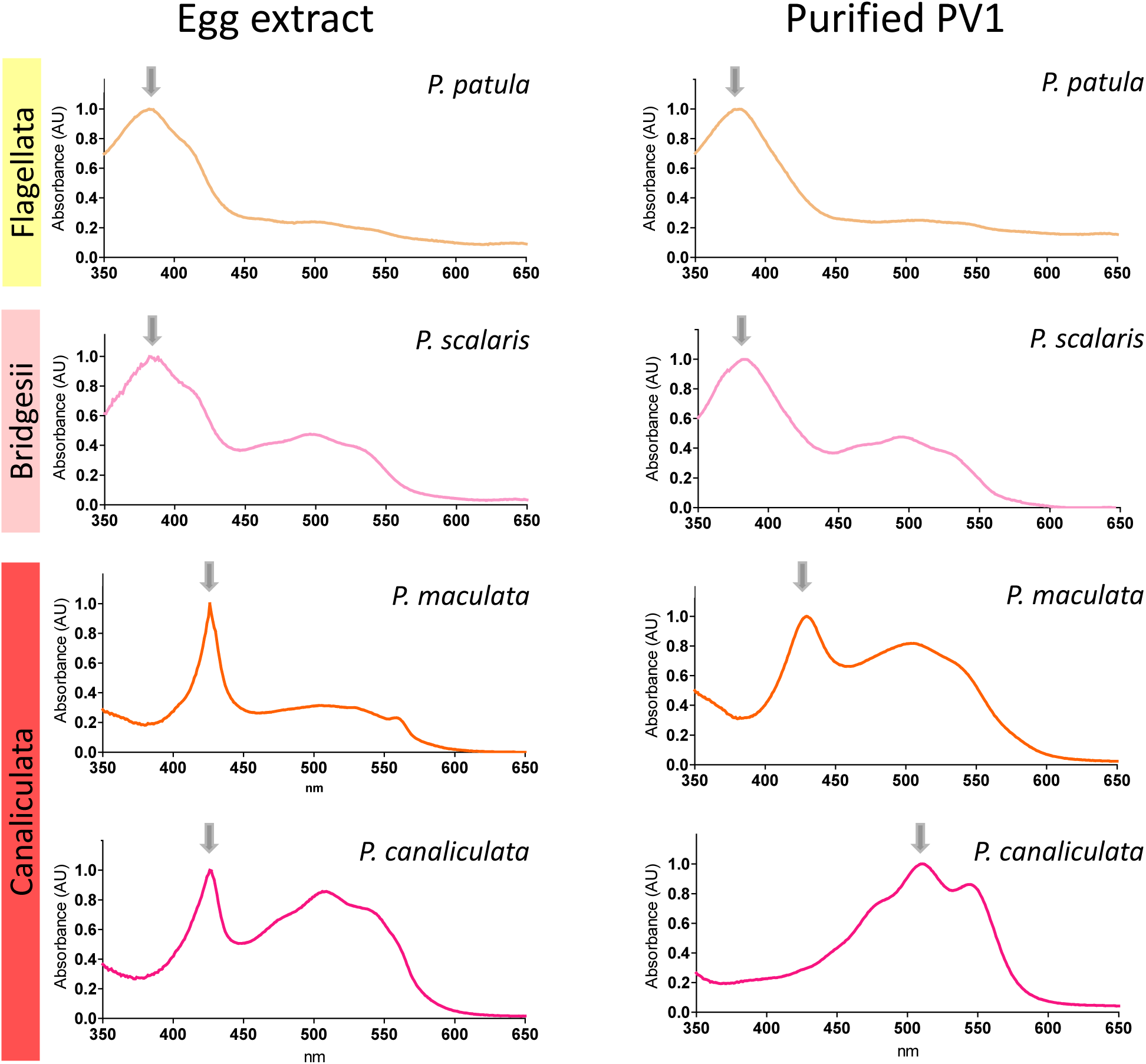
Pomacea eggs and PV1 absorption spectra shifts toward red in most derived clades. A. Egg extract (perivitelline fluid) B. purified egg carotenoproteins. Spectra are ordered from basal (top) to derived (bottom) clades. Arrows indicate the maximum of each spectrum to highlight the red-shift from basal to derived clades. Spectra were normalized for easy comparison. Data of P. scalaris egg carotenoprotein taken from Ituarte et. al. 2008.

### STRUCTURAL AND KINETIC STABILITY

#### Structural stability against pH, temperature, and chemical chaotropes

PpaPV1 remained stable in a wide range of pH. A slight alteration in the fine structure of the UV-visible spectrum (**Fig. S3A**) and an increase in fluorescence emission intensity (**Fig. S3B**) were only observed at pH=2.0. The absorption and emission spectra of PpaPV1 remained virtually unchanged even at temperatures of 80-85 °C (**Fig. S3 C-D**). Also, the electrophoretic behavior after boiling PpaPV1 for 60 min was unchanged, as reported for other *Pomacea* carotenoproteins (**Fig. S4**).

The chemical stability of PV1s showed an overall increase in fluorescence intensity and a systematic red shift of the spectra when increasing GdnHCl concentrations (**Fig. S5**). **Figure 3A** shows that the GndHCl unfolding transition of PpaPV1 reaches a plateau and experimental data fits a two-state model. The GdnHCl concentration required to reach the midpoint of the transition between both states (Cm) was lower in PmPV1 and PsSC than in PpaPV1 (the GdnHCl concentration to obtain fifty percent unfolded) (Figure 3B, Table 2). The disassembling/unfolding process followed by changes in the standard free energy ΔG°H_2_O, was higher in PpaPV1 than in PmPV1 (Figure 3B, Table 2) shows this parameter in different *Pomacea* spp. PV1s.

**Figure 3.**
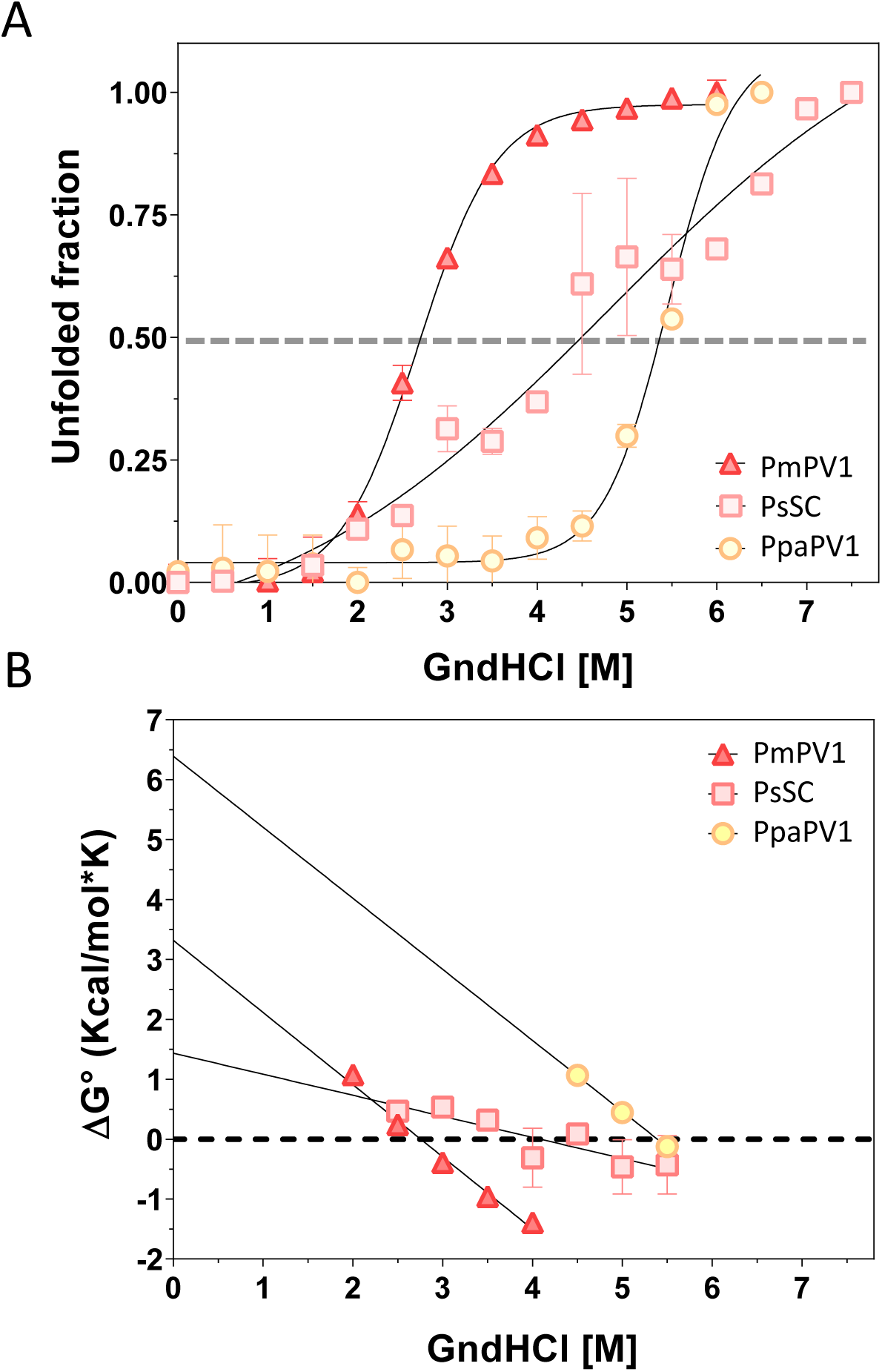
Structural stability of PV1 decreases in a clade-related fashion. Stability of PpaPV1, PsSC, and PmPV1 was evaluated by the unfold induced by GdnHCl. A. Unfolded population of PV1s in the equilibrium. B Dependence of the unfolding free energy (ΔG0) with GndHCl concentration. ΔG0H2O was calculated from the ordinate intercept. Cm: GndHCl concentration at ΔG0=0 (midpoint of the denaturing transition). PmPV1 data taken from Pasquevich et al. 2017.

**Table 2.** Thermodynamic parameters of Pomacea apple snail perivitellins unfolding induced by chemical treatment.

| Clade | Flagellata | Bridgesii |  | Canaliculata |  |
| --- | --- | --- | --- | --- | --- |
|  | *P. patula* | *P. scalaris* | *P. diffusa* | *P. maculata* | *P. canaliculata* |
| **PV1** | **PpaPV1** | **PsSC** | PdPV1<sup>£</sup> | **PmPV1** | PcOVO<sup>¥</sup> |
| **$\Delta G^0_{H_2O}$ (kcal/mol)** | **6.40 ± 0.32** | **1.43 ± 0.44** | 1.14 | **3.32 ± 0.19** | 3.48 ± 0.06 |
| **Cm (M)** | **5.4** | **4.1** | 3.4 | **2.8** | ND |

<sup>¥</sup> Data from Dreon et al. 2007; <sup>£</sup> Data from Brola et al. 2020; ND not determined; In bold: data from this study (See Material and Methods for experimental details).

#### Resistance to sodium dodecyl sulfate-induced denaturation

Proteins with a high energetic barrier between the folded and unfolded states are very resistant to unfolding and are considered kinetically stable. Comparison of the migration on polyacrylamide gels of identical boiled and unboiled PV1s previously incubated with SDS indicated PpaPV1 was resistant to SDS-induced denaturation. However, PV1s from Bridgesii and Canaliculata clades, display a partial loss and therefore some oligomers in the unheated samples disaggregates in their subunits (**Fig. 4A**).

**Figure 4.**
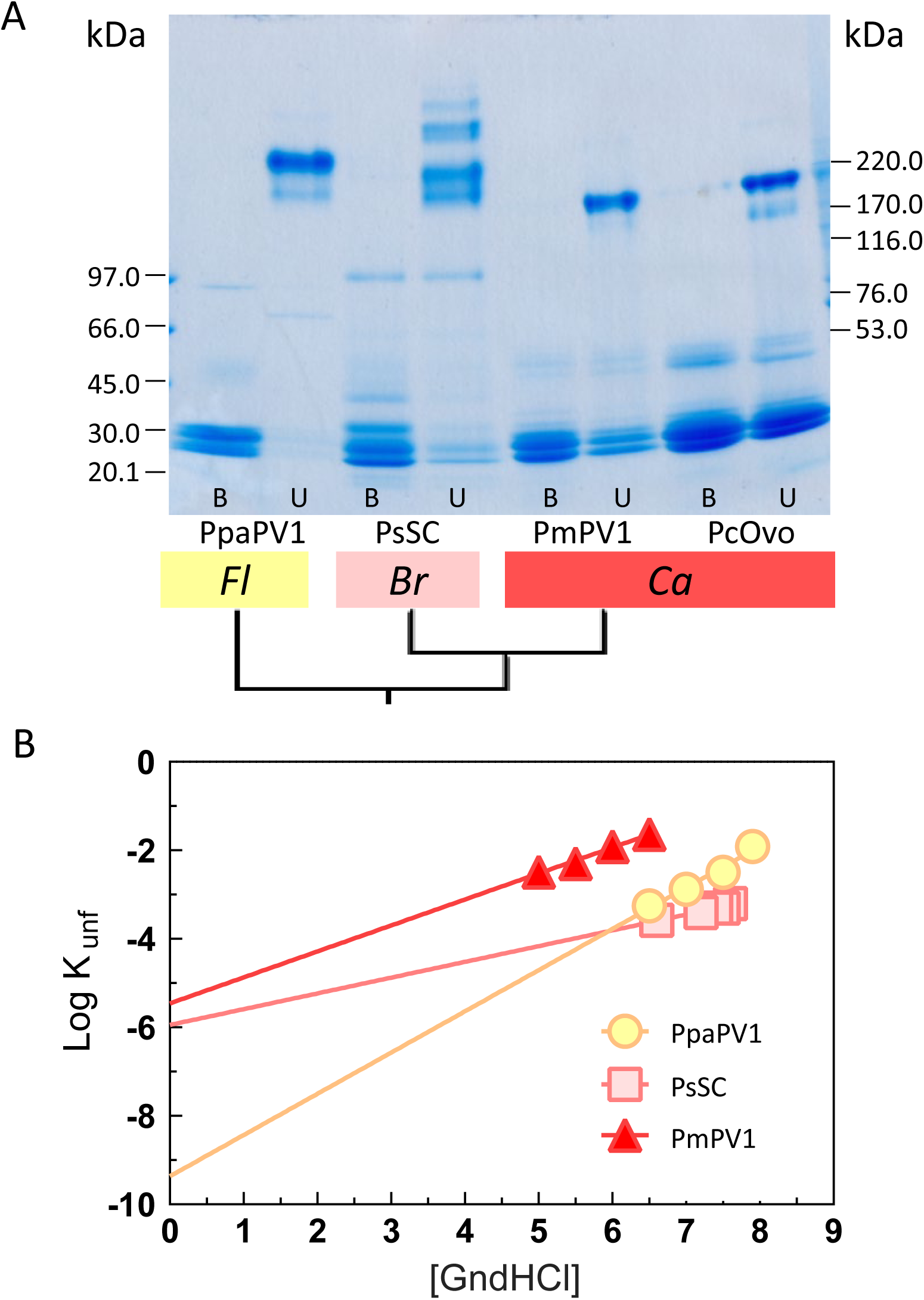
The PV1s of basal clades are hyperstable proteins extremely resistant to detergent treatment and chemical denaturation. A. SDS-PAGE of PV1s previously unheated (U) or boiled (B) in the presence of SDS detergent for 10□min and immediately loaded into the gel. PpaPV1 is more resistant to detergent treatment than those PV1s from other clades. A comparison with other hyperstable proteins in nature is given in Table 3. Fl: Flagellata; Br: Bridgesii; Ca: Canaliculata; B. Unfolding rates of PpaPV1 and PmPV1 under native-like conditions are shown by extrapolating the unfolding rate determined at different concentrations of GdnHCl to 0 M. The y-intercept of each extrapolation curve indicates the unfolding rate of the native protein. PpaPV1 has unfolding kinetics much slower than the orthologue of the most derived clade.

**Table 3.**
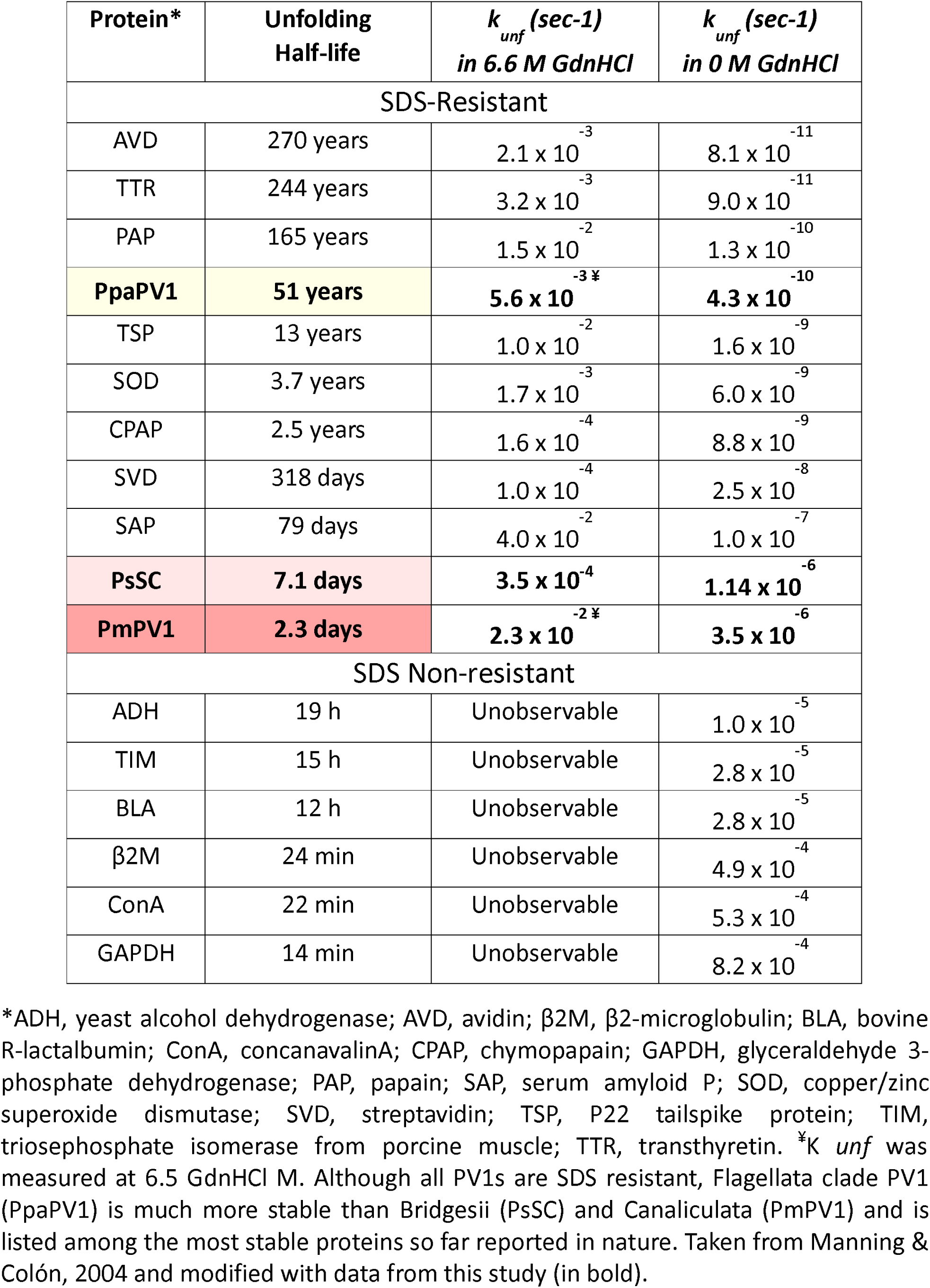
Comparison of the Half-Lives and Unfolding Rate Constants of snail PV1s and proteins resistant and not resistant to SDS. Proteins were sorted according to their half-life.

#### Unfolding rate and resistance to proteolysis: kinetic stability of PV1s

Subjecting carotenoprotein to increasing GdnHCl concentration and measuring the time they took to unfold allowed the calculation of the half-life of proteins in native conditions without the presence of the chaotrope. PV1s of the Flagellata, Bridgesii, and Canaliculata clades were assayed, namely PpaPV1, PsSC, and PmPV1. The rate at which PpaPV1 carotenoprotein unfolds is markedly lower than that of its derived clade counterparts. Consequently, the half-life of PpaPV1 was several orders greater than PsSC and PmPV1 (**Table 3**).

Limited proteolysis of PpaPV1 with Proteinase K, a fungal protease with broad specificity showed no evidence of PpaPV1 hydrolysis, while BSA (control) was completely digested (**Fig. S6**).

### FUNCTIONAL CHARACTERISTICS

#### Lectin activity

The carbohydrate-binding capacity of PpaPV1 evaluated by hemagglutination assays showed activity in a dose-dependent manner. The hemagglutinating activity was up to 0.16 μg/μL (**Figure 5A**). The specificity and relative affinity of PpaPV1 towards oligosaccharides structures were evaluated by a high-throughput glycan array assay. Albeit with low affinity, PpaPV1 showed a binding pattern to glycans related to the Blood A group containing a specific motif (GalNAca1-3(Fuca1-2)Galb1-4GlcNAc) (Figure 5B). The specificity toward oligosaccharides is shown in Table 4. Among them, the specificity of PpaPV1 with Blood group A type 2 antigens, as well as the lack of specificity toward sialic acid antigens, are remarkable.

**Figure 5.**
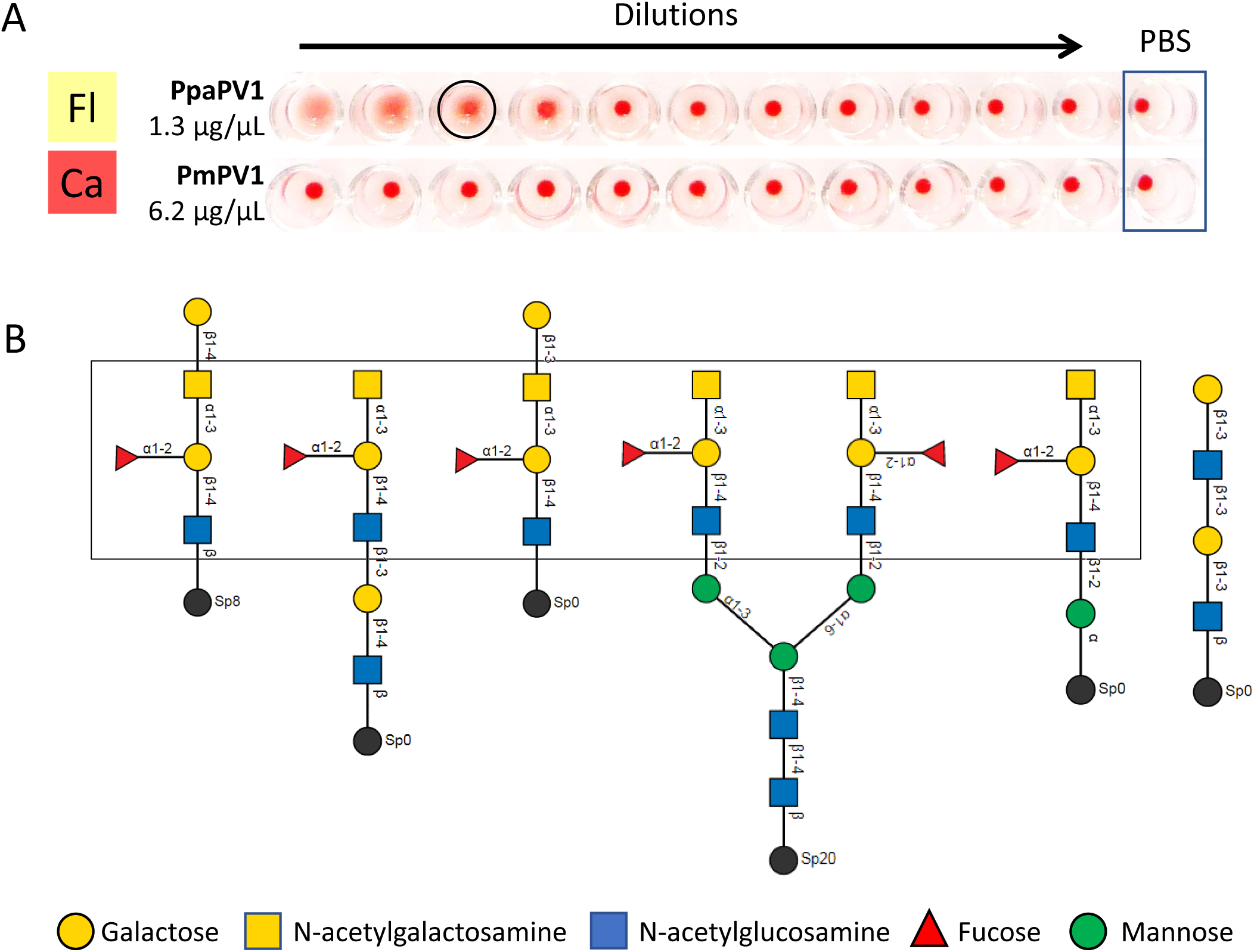
The Lectin capacity of Flagellata PpaPV1 is not as strong as Bridgesii PV1s and has narrower glycan binding motifs**. A.** Microplate well showing the hemagglutinating activity of *P. patula* purified PpaPV1 and PmPV1 from *P. maculata.* Circled well correspond to the last dilution that hemagglutinate. The last well of each row corresponds to PBS. PpaPV1 has moderate hemagglutinating activity while PV1 belonging to the derived canaliculata clade lacks this capacity. **B.** Main glycan structures recognized by PpaPV1 highlighting a common recognizing pattern: *GalNAc*a1-3*(Fuca1-2)Gal*b1-4*GlcNAc* (rectangle). The glycan structure plot was taken from the Consortium for Functional Glycomics (http://www.functionalglycomics.org). See Table 4 for more details.

**Table 4.** Main glycan structures recognized by PpaPV1.

| Rank | Oligosaccharide Structure | Average RFU | %CV |
| --- | --- | --- | --- |
| 1 | <u>GalNAca1-3(Fuca1-2)Galb1-4GlcNAcb1-2Mana1-6(GalNAca1-3(Fuca1-2)Galb1-4GlcNAcb1-2Mana1-3)Manb1-4GlcNAcb1-4GlcNAcb-Sp20</u> | 1081 ± 73 | 7 |
| 2 | <u>GalNAca1-3(Fuca1-2)Galb1-4GlcNAcb1-3Galb1-4GlcNAcb-Sp0</u> | 325 ± 40 | 12 |
| 3 | <u>Galb1-4GalNAca1-3(Fuca1-2)Galb1-4GlcNAcb-Sp8</u> | 318 ± 41 | 13 |
| 4* | <u>Galb1-3GlcNAcb1-3Galb1-3GlcNAcb-Sp0</u> | 309 ± 3 | 1 |
| 5 | <u>Galb1-3GalNAca1-3(Fuca1-2)Galb1-4GlcNAc-Sp0</u> | 239 ± 28 | 12 |
| 6 | <u>GalNAca1-3(Fuca1-2)Galb1-4GlcNAcb1-2Mana-Sp0</u> | 204 ± 20 | 10 |
| 7* | <u>(3S)Galb1-4(Fuca1-3)(6S)Glc-Sp0</u> | 163 ± 14 | 9 |

Binding intensities are expressed as the mean of relative fluorescence units (RFU) ± 1SD, N=4. %CV = 100 x SD. Full data of PpaPV1 glycan specificity is available as Supporting Information. \*not PpaPV1 concentration-dependent binding.

#### *In vitro* simulated gastrointestinal digestion

PpaPV1 resists hydrolysis when exposed sequentially to 2 h of gastric and duodenal phases (Fig. 7S) while BSA (control) was readily degraded. PpaPV1 maintained its electrophoretic behaviour for up to 120 min.

## DISCUSSION

### The complexity of the heteroligomerization increases in the most derived species

PV1s heteroligomers are composed of combinations of related subunits, probably paralogues that arise by duplication and speciation from an orphan gene (Sun et al. 2012). Based on the phylogenetic hypothesis proposed by Hayes et al. (2009) analysis of PV1 from the most basal clade of the genus (Hayes et al. 2009) allowed for the first time to investigate the evolution of structure, stability, and functional features of proteins within a single genus. Figure 6 synthesizes the hypothesis of PV1 evolution from our current knowledge of this. The first conclusion is that while the mass and size of PV1 particles remained similar along evolution and their subunits have roughly the same molecular weight regardless of the phylogenetic position, marked structural changes are present. Particles seem to have acquired different post-translational modifications and surface structural changes were further evidenced by the lack of cross-reactivity of Flagelatta and Bridgesii PV1 with antibodies directed against Canalicula PV1s. As expected from the orphan gene origin of this novel family of proteins, no similarities with sequences reported outside ampullarids were observed in PpaPV1. Besides, while oligomers from derived clades combine 6 different paralogue subunits, PpaPV1 combines only 3 different subunits, one with no similarities with any other PV1s, further suggesting that subunits aroused by gene duplication (Sun et al. 2012, Pasquevich et al. 2017, Ip et al. 2019, Brola et al. 2020) and some subunits were lost along evolution. In addition, the Flagellata and Bridgesii clades carotenoproteins share common spectroscopic features that change markedly in PV1s from the most derived species.

**Figure 6.**
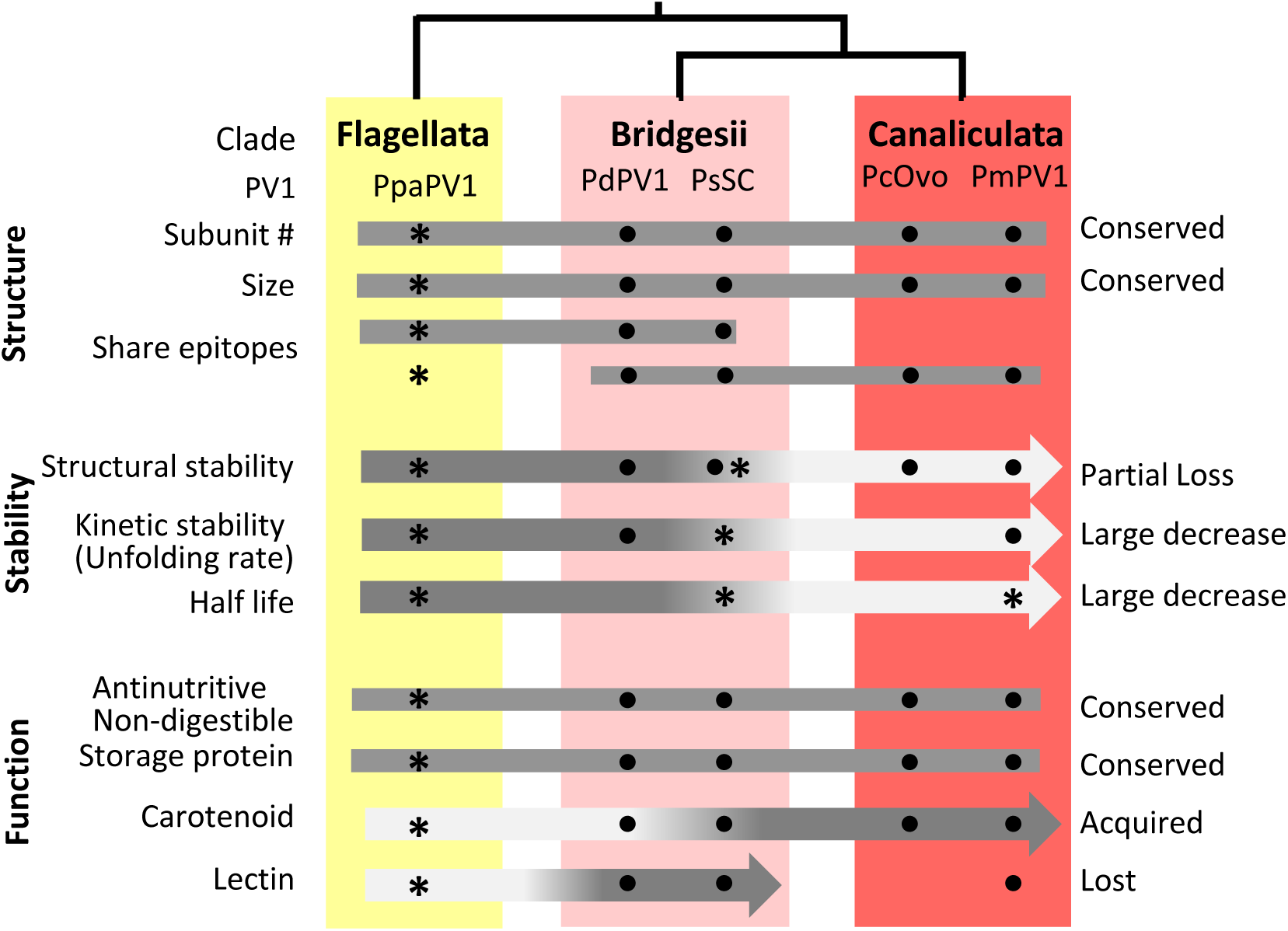
Hypothesis of the evolution of structure, stability and functional features of PV1 carotenoproteins in *Pomacea* genus. *Data from this study. Dots indicate proteins studied for the trait. *Pomacea* phylogeny was based on Hayes *et al*. 2009. Data was taken from Brola et al. 2020; Dreon et al. 2004a, 2004b; Ituarte *et. al.* 2008, 2010, 2012; Pasquevich et al., 2014, 2017.

### Evolutionary significance of the stability loss of apple snail eggs carotenoproteins

The term *protein stability* indicates the ability to retain the native conformation when subjected to physical or chemical manipulation. PpaPV1 is structurally highly stable, rather difficult to unfold (denature) by near-boiling temperatures, extreme pHs and high concentrations of a denaturing chemical, a common feature shared with Bridgesii and Canaliculata carotenoproteins (Ituarte et al., 2012, Dreon et al., 2007, Dreon et al., 2007, Pasquevich et al., 2017, Brola et al. 2020). However, PpaPV1 displayed a much higher resistance to chemical denaturation than the other PV1s.

Our results strengthen the notion that long half-live proteins (kinetically stables) withstand SDS detergent retaining a rigid folded core that only unfolds when boiled (Manning and Colon 2004). In our study, PpaPV1 shows a much longer half-life and a greater resistance to denaturation by SDS than other PV1s. Moreover, PpaPV1 is among the most stable proteins so far reported while PmPV1 lies in the transition between the most stable proteins and those that are not (Table 2), even considering these values may have errors > 25% (Manning & Colón, 2014). PpaPV1, as other *Pomacea* spp. carotenoproteins are also exceptionally resistant to proteolysis, suggesting that a common mechanism may account for their resistance to SDS and proteolytic cleavage (Manning & Colón, 2004). More broadly, kinetically stable proteins are typically extracellular and several protecting from oxidative stress have evolved as kinetically stable (Colón et al., 2017). In this regard, PV1s have both features as they are located in the fluid surrounding the embryos and their pigments provide antioxidant protection to eggs (Dreon et al., 2004a). Along the evolution of PV1s we noticed a partial loss of stability, but only to a certain point as the removal of carotenoids does not affect the stability of PcOvo (Dreon et al., 2007). This supports the idea that PcOvo stability and probably other members of the PV1 family favours its physiological role in the storage, transport, and protection of carotenoids during snail embryogenesis.

The hyperstability of PpaPV1 basal carotenoprotein could have allowed tolerance to mutations, *i.e.*, acquiring new functions (functional evolution) without losing the native structure, in agreement with Bloom *et al*. (2006), which proposed that the high stability allows the evolvability of proteins.

The high kinetic stability of these orthologs defense proteins is a vital property of their protective role and is related to snail fitness (reproductive success).

### Roles of PpaPV1 and evolutionary functions of PV1s

Terrestrial egg deposition in *Pomacea* was a key adaptation to avoiding aquatic predation and/or parasitism (Sun et al., 2019). This evolutionary driver modulated snail egg defenses and under this selective pressure, PV1 orthologous, while retaining some roles, underwent major functional adaptations. Thus, PV1s kept their ancestral traits as storage proteins to nurture embryos but not digestible by predators, whereas gradually lost their stability and gained new roles (**Figure 6**) a known tradeoff between the evolution of new-function and protein stability (Tokuriki & Tawfik, 2009b). The unfolding speed of PV1 in the basal clade is dramatically lower than that of the most derived clade homologue and we can hypothesize that this loss of stability allowed in Canaliculata proteins structural changes favoring a better stabilization of more polar carotenoids and the ability to accommodate larger quantities of this pigment. This agrees with other studies that indicate the loss of stability could have contributed to the gain of new functions (Bloom et al. 2006). This PV1 feature came at the price of losing the lectin capacity as it is discussed below.

To the best of our knowledge, PV1 evolution is one of the few examples taken from nature where the tradeoff between the stability and evolvability of a protein is reported. One interesting aspect is the the loss of the lectin capacity in the most derived clade. This is at first intriguing considering its significant role in the defenses against predation (Brola et al., 2020) but may be explained by the evolutionary novelty in Canaliculata clade of a dual enterotoxic/neurotoxin lectin (PV2) combining two ancient immune proteins (Giglio et al., 2020). On the other hand, the Bridgesii clade not only retained Flagelatta lectin capacity, but PpaPV1 moderate lectin activity gives way to PV1s with higher affinity to glycans and a broader specificity indicative of at least two high-affinity recognition sites in *Bridgesii* clade (Ituarte et al., 2018). Remarkably, PsSC and PdPV1 binding recognition patterns include Galβ1-3GalNAc and a common sialic acid in vertebrate gangliosides (Brola et al., 2020; Ituarte et al., 2018) that are virtually absent in the glycans patterns recognized by PpaPV1. The inability of PpaPV1 to recognize gangliosides, and the nearly limited group A type II antigen recognition patterns, suggest that a single recognition site would be present in this ancient PV1. It can be speculated that the partial loss of PV1s stability in Bridgesii may have favored the evolution of their improved glycan recognition and binding strength capabilities.

## CONCLUSION

This study shows the phylogenetic differences of the structural and functional features of a novel family of invertebrate egg proteins originating from orphan genes. The basal clade contains one of the most kinetically stable proteins known to date. More broadly, and supported by the currently accepted evolutionary hypothesis of the genus phylogeny, this study provides one of the few examples taken from nature, as opposed to directed evolution in the laboratory, showing that during orthologue evolution there is a tradeoff between a loss of structural and kinetic stability and a simultaneous acquisition of new defensive traits. The extent to which these mechanisms are evolutionary steps or alternative trajectories in the evolution of selective expression of defensive strategies in the eggs is an open question. This work also increases the knowledge of ampullarids biology referring to the evolution of the complex defense system of *Pomacea* apple snail eggs.

Considering that *Pomacea* has split from its sister genus just about 28 mya (Sun et al., 2019), the study also highlights how a rapid evolution of structure-function features of reproductive proteins accompanied the spread and diversification of *Pomacea* across freshwater habitats. Predator-induced protein evolution may have contributed to the evolved defence strategies and may have contributed to canaliculata snails’ worldwide invasiveness.

## ACKNOWLEDGMENTS

MYP and HH are members of CONICET, Argentina. MSD is a member of CICBA, Argentina. We thank L. Bauzá for her help in PpaPV1 purification and Dr. L. Falomir-Lockard for help in DLS analysis. We acknowledge the participation of the Protein-*Glycan* Interaction Resource of the CFG and the National Center for Functional Glycomics (NCFG) at Beth Israel Deaconess Medical Center, Harvard Medical School (*supporting grant R24 GM137763*).

